# The effects of chronic neuropathic pain on the self-administration of highly potent MOR agonist, fentanyl

**DOI:** 10.1101/2024.11.19.624389

**Authors:** Gwendolyn E. Burgess, John R. Traynor, Emily M. Jutkiewicz

## Abstract

There is significant overlap between chronic pain and opioid use disorder (OUD) patient populations such that approximately 50-65% of chronic pain patients have OUD. However, we understand relatively little about how chronic, long-lasting pain states alter ongoing self-administration of opioid analgesics. Thus, the goal of this study was to determine if chronic neuropathic pain altered the ongoing self-administration of fentanyl, or a non-opioid drug of abuse, cocaine. Animals were trained to self-administer fentanyl or cocaine in a multi-dose self-administration procedure composed of five 25-min components, exposing animals to multiple doses of drug per day. Operant behavior was established prior to induction of chronic pain via the spared nerve injury (SNI). Animals were allowed 72 hours of post-operative recovery and resumed self-administration on post-operative day 4. All animals dose-dependently self-administered fentanyl prior to surgery. On post-operative day 4, both sham and SNI groups showed a significant decrease in fentanyl self-administration. By post-operative day 9, fentanyl intake was no longer significantly different from pre-surgical intake. Over the course of 4 weeks of self-administration, there was an increase in intake of specifically the 10 ug/kg/inf dose of fentanyl. Cocaine self-administration was not altered at any point following either surgery. Collectively, these results suggest that SNI-induced hypersensitivity failed to alter the reinforcing effects of fentanyl, or non-opioid drug of abuse, cocaine. Future studies should evaluate the abuse potential of lower efficacy MOR agonists such as nalbuphine or buprenorphine, as small changes were observed in fentanyl-maintained behavior over time in both SNI and sham groups.

**Significance statement:** MOR agonists are excellent analgesics; however, they are not first-line treatments for chronic pain, in part due to abuse potential. This study demonstrates that small, significant rightward shifts in the fentanyl dose response curve were observed following induction of *both* sham and spared nerve injury (SNI) (chronic neuropathic pain-like) states, suggesting these changes were observed independent of pain state. These data indicate that SNI-induced neuropathic pain failed to alter the ongoing self-administration of highly potent MOR agonist, fentanyl.

## Introduction

Both chronic pain and opioid use disorder (OUD) are substantial public health concerns with overlapping patient populations, such that 50-65% of chronic pain patients have been diagnosed with OUD (Hser et al., 2017; Latif et al., 2021). While mu opioid receptor (MOR) agonists are useful analgesics for moderate to severe pain, they are rarely considered first-line treatments for chronic pain because repeated use of MOR agonists leads to the development of tolerance and requires escalation of dose, physical dependence, and abuse liability. This is particularly concerning since many (∼69%) patients with chronic neuropathic pain are treated with MOR agonists at some point for pain management, and ∼20% of chronic pain patients are treated with opioids persistently/frequently (Hoffman et al., 2017). While exposure to and use of MOR agonists increases abuse liability, some studies have also suggested that pain itself a risk factor for the development of OUD (Bauman et al., 2023; Brummet et al., 2017; Katz et al., 2013). However, we know relatively little about whether or not chronic pain alters the reinforcing effects of MOR agonists.

Previous preclinical studies have examined the rewarding or reinforcing effects of MOR agonists in the presence of various pain states, but the results have been heterogeneous. Some studies show an *increase* in the rewarding (Cahil et al., 2013; Lim et al., 2014; Navratilova et al., 2020; Zhang et al., 2014) or reinforcing effects (Colpaert et al., 2001; Hipólito et al., 2015; Higginbotham et al., 2022) of MOR agonists in the presence of a pain state, while other pre-clinical experiments have shown a *decrease* in rewarding or reinforcing effects of opioids (Ozaki et al., 2002; Suzuki et al., 1996; Nazarian et al., 2021; Narita et al., 2005; Wade et al., 2003; Martin et al., 2007). Further, other studies showed *no impact* of pain states on opioid induced reinforcing effects (Barattini et al., 2023; Hou et al., 2015; Reiner et al., 2021) or opioid rewarding effects (Shippenberg et al., 1988). In previous self-administration studies specifically, there are many methodological differences; however, one difference worth considering is the duration of the pain state used. Many previous studies have utilized acute or sub-chronic pain states (Baratinni et al., 2023; Hippolito et al., 2015; Higginbotham et al., 2022; Hou et al., 2015; Reiner et al., 2021; Wade et al., 2003), which prevents the long-term evaluation of MOR agonist-maintained behavior.

In addition, only three studies have assessed if pain states alter established, ongoing opioid self-administration (Baratinni et al, 2023; Neelakantan et al., 2017, Reiner et al., 2021). Two of these studies utilized short acting (lactic acid, capsaicin; Reiner et al., 2021) pain states or sub-chronic pain states (CFA; Baratinni et al., 2003), and both of these studies found no effect of pain state on MOR agonist-maintained behavior. The third used paclitaxel-induced neuropathic pain and investigated if developing neuropathic pain altered breakpoint responding for morphine infusions in mice (Neelakantan et al., 2017). This study found that the injured animals had consistent, stable breakpoint responding for morphine, while the saline-treated group displayed a decrease over time. However, no study has examined the impact of a fully developed, chronic neuropathic pain state on the ongoing self-administration of opioids. Therefore, the goal of this study was to examine (within-subject) the reinforcing effects of fentanyl before and after the induction of a persistent, long-lasting pain state caused by the spared nerve injury (SNI).

## Methods

### Animals

All rats were at least 8 weeks old at the beginning of the study. Rats were ordered from Envigo labs (Indianapolis, IN). Following delivery, rats were given one week to habituate to the animal housing facility. All rats were housed in standard cages with corncob bedding. Water was available ad libitum. The housing facility was kept on a 12-hr light dark cycle, and all experiments were conducted in the light cycle.

Following intravenous catheter implant, rats were single housed for the remainder of the study of the study. These rats were food restricted 24 hours prior to the start of self-administration experiments. Males were fed 5 pellets and females were fed 4 pellets of standard rat chow (5LOD, LabDiet) to maintain rats at approximately 90% of their free-feeding body weight. Following SNI or sham surgery, rats were given wooden gnawing blocks and enviropacks as enrichment.

For the Randall Selitto assays, rats were single housed for 7-10 days following surgery (femoral catheter implant & SNI or sham surgery performed on the same day) until sutures were removed, then were group housed for the remainder of the study with food available ad libitum. Following surgery, rats were given wooden gnawing blocks and enviropacks as enrichment.

### Surgery

#### Anesthesia

For all surgeries, rats were deeply anesthetized with 90 mg/kg ketamine and 10 mg/kg xylazine, injected intraperitonially (i.p.). Rats were given carprofen (5 mg/kg) subcutaneously (s.c.) prior to surgery as well as 24-hr later for pre- and post-operative analgesia. Following SNI or sham surgery, rats were given 5 mg/kg carprofen s.c. 24- and 48-hr after surgery.

#### Femoral Vein Catheter and Backplate Implant

Rats were implanted with a chronic, indwelling intravenous catheter. Briefly, the surgical incision sites were shaved, cleaned, and sterilized. An incision was made in the left lower abdomen ∼1 cm from the left leg and the femoral vein was exposed. A catheter (Micro-renathane tubing MRE-033-male; MRE-040-female, Braintree Scientific Inc., Braintree, MA) was inserted and passed subcutaneously to the scapulae where it was connected to the backplate (PlasticsOne, Roanoke, Virginia, USA, 8I313000BM14). The mesh of the backplate was sutured to the muscle, and then the skin closed around the backplate.

#### Spared Nerve Injury or Sham surgery

These surgeries were carried out as described in Decosterd & Woolf (2000). Briefly, the left femoral muscle was opened to expose the sciatic nerve. A 2 mm section of the peroneal and tibial nerves were removed, while the sural nerve was not disturbed. The exposed ends of the peroneal and tibial nerves were sutured to the underlying muscle to prevent regrowth. The sham surgery consisted of only the muscle incision, with no manipulation to any branch of the sciatic nerve. In both SNI and sham surgeries, the muscle was closed with absorbable suture, and the skin was closed with surgical staples (9mm; Braintree Scientific, Braintree, MA).

### Apparatus

Self-administration experiments were conducted in 18 standard MED PC operant chambers (ENV-008CT, Med Associates, St. Albans, VT), which were housed in sound-attenuating chambers (ENV-018CT). These chambers were equipped with two nosepoke manipulandum on the right side of the chamber. A house light (white, ENV-114BM) was located on the top of the left side of the chamber. A variable speed pump (PHM-107) was used to deliver drug infusions. A 20 mL syringe was connected to the single channel, luer lock swivel (375/22PLS, Instech, Plymouth Meeting, PA) by luer lock tubing. Tygon tubing (100-80) was used to connect the swivel to the backplate, and the tygon tubing was contained in a stainless-steel spring. Data were collected via MED-PC software (SOF-735).

### Procedures

#### Randall Selitto & Testing of Mechanical Hypersensitivity

To evaluate mechanical thresholds, a paw pressure applicator (Randall Selitto, IITC life sciences, Woodland hills CA) was used to apply continuously increasing amounts of pressure to the contralateral (non-surgerized limb) or ipsilateral (injured, sham injured) paws until the pressure caused a retraction of the hindpaw. The pressure resulting in hind paw retraction was recorded from the Analgesiometer with a 300 g pressure used as the maximum cutoff.

Prior to testing, rats were habituated to the hammock-like restraint over three days. On day one, rats were placed into the restraint for 5 min, and the habituation time was increased by 5 min/day until 15 min of habituation was achieved on the third day.

For all experiments, drugs were given i.v. and catheters were flushed with 0.5 mL heparinized saline daily for the duration of the experiment to maintain patency.

#### Self-Administration

##### Single component training sessions

Rats in self-administration experiments were allowed 7-10 days of post-operative recovery following catheter implant. All catheters were flushed daily during recovery with 0.5 mL 50 usp/mL heparinized saline. All self-administration sessions started with a catheter pre-fill infusion of (50 uL; confirm value). Rats were trained to self-administer fentanyl (0.0032 mg/kg/infusion) or cocaine (0.32 mg/kg/infusion) in daily 90 or 60 min sessions, respectively. Responding on the active nosepoke (illuminated) under a fixed ratio (FR)1 schedule of reinforcement resulted in delivery of drug, while responding on the inactive nosepoke had no scheduled consequences. After each infusion of drug, there was a 10 second time-out. The work requirement increased, across days, to a FR5 schedule of reinforcement. To proceed to the multi-dose procedure (details below), 5 consecutive days of stable responding were required. Stable responding was defined as no more than a 30% deviation in number of active responses (with no upward or downward trends) between days and <30% total responses occurred on the inactive nosepoke.

##### Multi-dose procedure

A multi-component, daily self-administration task was used to evaluate a range of doses of fentanyl *or* cocaine. Each session consisted of five, 25-min components with two-min blackouts between components during which responding had no scheduled consequence. For cocaine sessions: 0 (no infusion), 0.032, 0.1, 0.32, and 0.56 mg/kg cocaine were used across the components. For fentanyl sessions: 0 (no infusion), 0.32, 1, 3.2, 10, and 32 *u*g/kg/inf fentanyl were tested in overlapping dose response curves. Doses were presented in a semi-random order such that the first component was always 0 mg/kg (no infusion) and the last component was always the largest dose tested (cocaine-0.56 mg/kg/inf; fentanyl 10 or 32 *u*g/kg/inf). Components 2, 3, and 4 were randomized daily. Following 5 days of stable responding (same criteria as above), rats underwent either SNI or sham surgery. Following SNI or sham surgery, rats were allowed three days of post-operative recovery. Rats resumed self-administration on post-operative day four.

### Drugs

Fentanyl citrate was obtained from Sigma Aldrich (Burlington, MA). Cocaine hydrochloride was obtained from NIDA Drug Supply. Ketamine hydrochloride was obtained from Hospira, INC (Lake Forest, IL). Xylazine was obtained from Hospira (Lake Forest, IL). Carprofen (Rimadyl) was obtained from (Zoetis, Parsippany, NJ). Fentanyl and cocaine were dissolved in sterile, physiological saline and delivered intravenously (i.v.). Xylazine was diluted from the stock solution to 20 mg/mL in sterile water. Carprofen was diluted to 5 mg/mL in sterile saline. Heparin was diluted to 50 usp/mL in sterile saline to produce heparinized saline. Xylazine and ketamine were delivered intraperitonially and carprofen was delivered subcutaneously.

### Statistical Analysis

Data were analyzed by 3-way (dose, surgical status, time OR dose, surgical status, sex) or 4-way (dose, time, surgical status, sex) repeated measure ANOVAs in SPSS version 29. Significant main effects or interactions were followed up with one-way or two-way ANOVA’s and post hoc analyses. Only main effects or interactions necessary for interpretation are reported. Corrections were made, if necessary, according to Mauchly’s W criteria.

We intended to consider sex as a biological variable in all experiments. Unfortunately, we encountered difficulties such that we were unable to maintain catheter patency until the end of the experiment in female rats. Therefore, data are underpowered to detect sex differences. For Randall Selitto experiments, catheter patency was only needed for 9 days, and we were able to include sex as a biological variable.

Data in the graphs represent only animals that finished the procedure or at least progressed to 2 weeks following surgery.

#### Binned Data

For self-administration data, pre-surgical dose response curves were plotted as an average of the 5 stable days prior to surgery. Day four and day 9 dose effect curves are dose response curves from individual days. SNI/sham 2-week dose response curves reflect day 10-14 (days 4-9 removed for above analysis). SNI/sham 4-week dose response curves reflect days 24-28 (last 5 days of experiment) and this abbreviated time window is shown to match the other timepoints.

## Results

### Figure 1. Fentanyl self-administration prior to surgery and on post-operative days 4 and 9

Figure 1A and D show fentanyl-maintained responses for each dose of fentanyl on post-operative days 4 and 9 in sham and SNI groups. There was a significant main effect of dose [F(2.23, 31.51)=29.16, p<0.001, η^2^_p_ =0.68], suggesting that fentanyl maintained responding in a dose-dependent manner. Surprisingly, there was no main effect of time [F(2, 28)=1.69, p=0.20, η^2^_p_ =0.11]. However, there was a significant interaction between dose and time [F(2.96, 41.50)=0.52, p<0.001, η^2^_p_ =0.65], suggesting that fentanyl-maintained responding shifted over time. This was likely due to the noticeable decrease in responding on day 4 in both sham and SNI groups as well as small differences in responding for 1 μg/kg fentanyl on day 9 as compared with pre-surgical dose effect curves.

**Figure 1.**
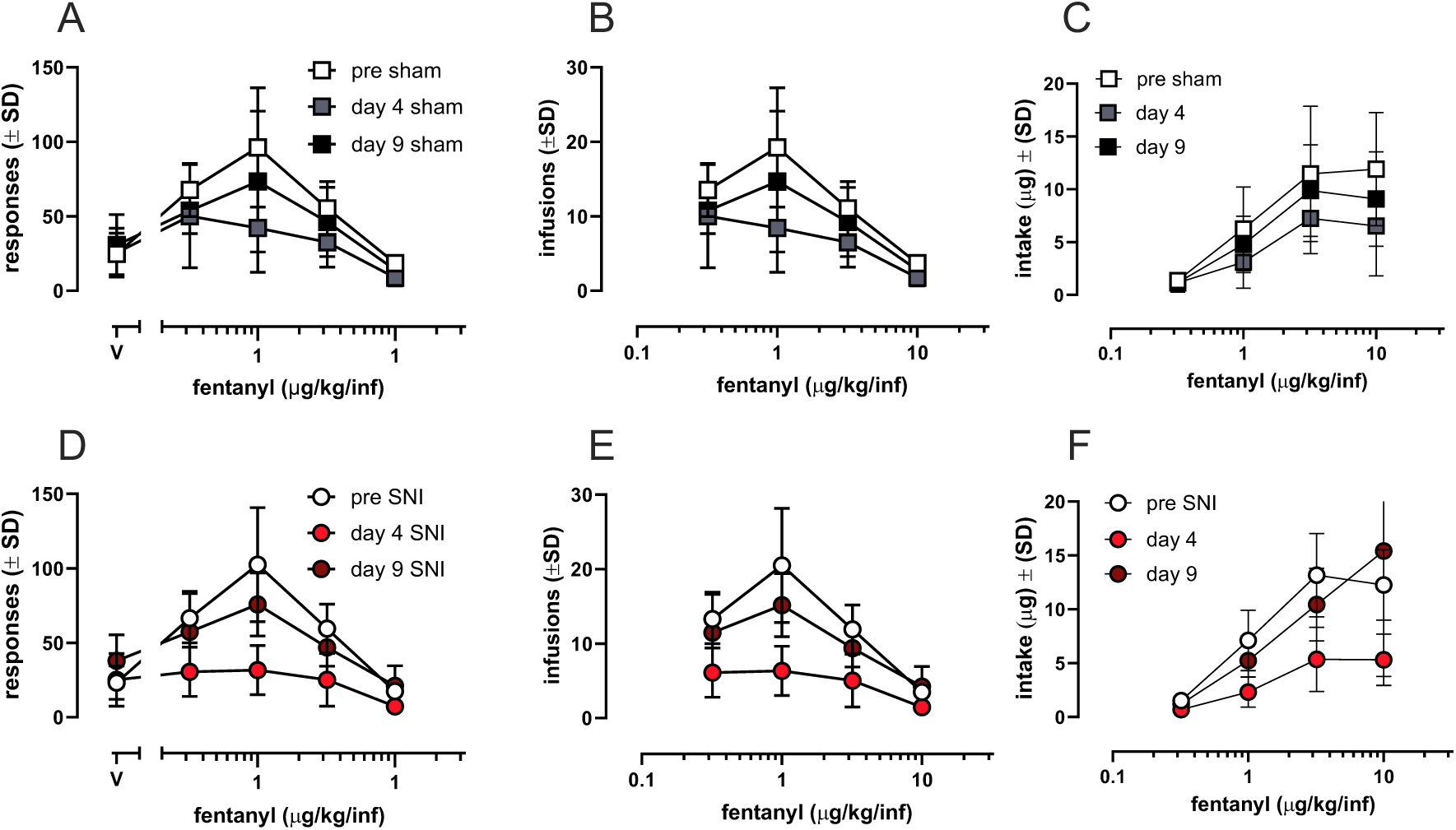
Self-administration of fentanyl prior to surgery and on post-operative days 4, 9 Data are plotted as responses for fentanyl (A, D), infusions of fentanyl earned (B, E), and total intake (μg) of fentanyl (C, F). Fentanyl maintained self-administration in all rats under a fixed ratio (FR)5 schedule of reinforcement prior to surgery. On post-operative day 4, there was a significant reduction in fentanyl self-administration in both sham (A-C) and SNI (D-F) groups; however, by day 9 following surgery there was no significant difference in fentanyl-maintained responding compared to pre-surgical dose effect curves. Data are plotted as the mean ± SD. Each data point represents 8-10 rats and include both males and females.

There was no main effect: surgical status [F(1, 14)=0.23, p=0.64, η^2^_p_ =0.02]. There were some changes in fentanyl-maintained responding; however, these changes are not due to surgical status [interaction: time* surgical status, F(2, 28)=0.099, p=0.91, η^2^_p_ =0.007]. Further, there was no significant three-way interaction between time, dose, and surgical status [F(2.96, 41.50)=0.52, p=0.67, η^2^_p_ =0.04]. Together, these data suggest that, although fentanyl-maintained responding was altered over time, surgical status did not explain these changes.

Figure 1B and E show the number of infusions of fentanyl earned for each dose of fentanyl on post-operative days 4 and 9. There was a significant main effect of dose [F(1.94, 27.14)=75.05, p<0.001, η^2^_p_ =0.84], suggesting different numbers of infusions were earned per dose. The number of fentanyl infusions earned was altered over time, reflected by a significant main effect of time [F(2,28)=20.27, p<0.001, η^2^_p_ =0.59]. Further, there was a significant interaction between dose and time [F(6, 84)=4.40, p<0.001, η^2^_p_ =0.24], which likely reflects that the number of infusions earned was most different at lower doses of fentanyl.

Though fewer infusions of fentanyl were earned on day 4 post-surgery, this change was not explained by surgical status, supported by no main effect of surgical status [F(1, 14)=0.23, p=0.64, η^2^_p_ =0.02] and no interaction between time and surgical status [F(1, 14)=0.23, p=0.64, η^2^_p_ =0.02]. Additionally, there was no three way interaction between time, dose, and surgical status [F(6, 84)=0.19, p=0.98, η^2^_p_ =0.01]. Collectively, these data suggest that the decrease in fentanyl infusions earned on post-operative day 4 was not explained by surgical status.

Figure 1C and E show the total fentanyl intake for each dose of fentanyl on post-operative days 4 and 9. Fentanyl intake was dose-dependent, reflected by a main effect of dose [F(1.93, 26.98)=103.90, p<0.001, η^2^_p_ =0.88]. Fentanyl intake was altered over time, reflected by a main effect of time [F(2, 28)=16.22, p<0.001, η^2^_p_ =0.54]. This was further supported by a significant time by dose interaction [F(2.21, 84)=3.88, p=0.03, η^2^_p_ =0.22], which likely represents the decrease in fentanyl intake on post-operative day 4.

Though fentanyl intake was subject to changes over time, these changes were not explained by surgical status, supported by no main effect of surgical status [F(1, 14)=0, p=0.998, η^2^_p_ =0] as well as no interaction of time and surgical status [F(2, 28)=2.05, p=0.15, η^2^_p_ =0.13]. Lastly, there was no three-way interaction of time, surgical status, and dose [F(6, 84)=1.40, p=0.23, η^2^_p_ =0.09]. Collectively, these results suggest that the decrease in fentanyl intake on post-operative day 4 was not explained by surgical status.

### Figure 2. Fentanyl self-administration prior to surgery and over 2- to 4-weeks following surgery

Figure 2A and D show the fentanyl-maintained responding for each dose of fentanyl 2- or 4-weeks following surgery. Fentanyl maintained dose dependent responding, reflected by a main effect of dose [F(2.15, 25.85)=71.05, p<0.001, η^2^_p_ =0.86]. There was no difference in fentanyl-maintained responding in SNI and sham groups, reflected by no main effect of surgical status [F(1, 12)=3.68, p=0.08, η^2^_p_ =0.24] (Fig. 2A, D). There was no main effect of time such that fentanyl-maintained responding did not significantly change from pre- to post-surgical timepoints [F(2, 20)=2.44, p-0.11, η^2^_p_ =0.20]. However, there was an interaction between dose and time [F(8, 96)=2.40, p=0.02, η^2^_p_ =0.17], suggesting that there were small, but significant, changes in responding for individual doses over time. For example, responding for 10 *u*g/kg/inf fentanyl at 4 weeks post SNI was increased as compared with pre-surgery in both sham and SNI groups. Additionally, responding for 1 *u*g/kg/inf decreased 2- or 4-weeks post SNI as compared with pre-surgery. However, there was no three-way interaction between surgical status, dose, and time [F(8, 96)=1.08, p=0.38, η^2^_p_ =0.083]. All other interaction terms failed to reach significance (p’s>0.12).

**Figure 2.**
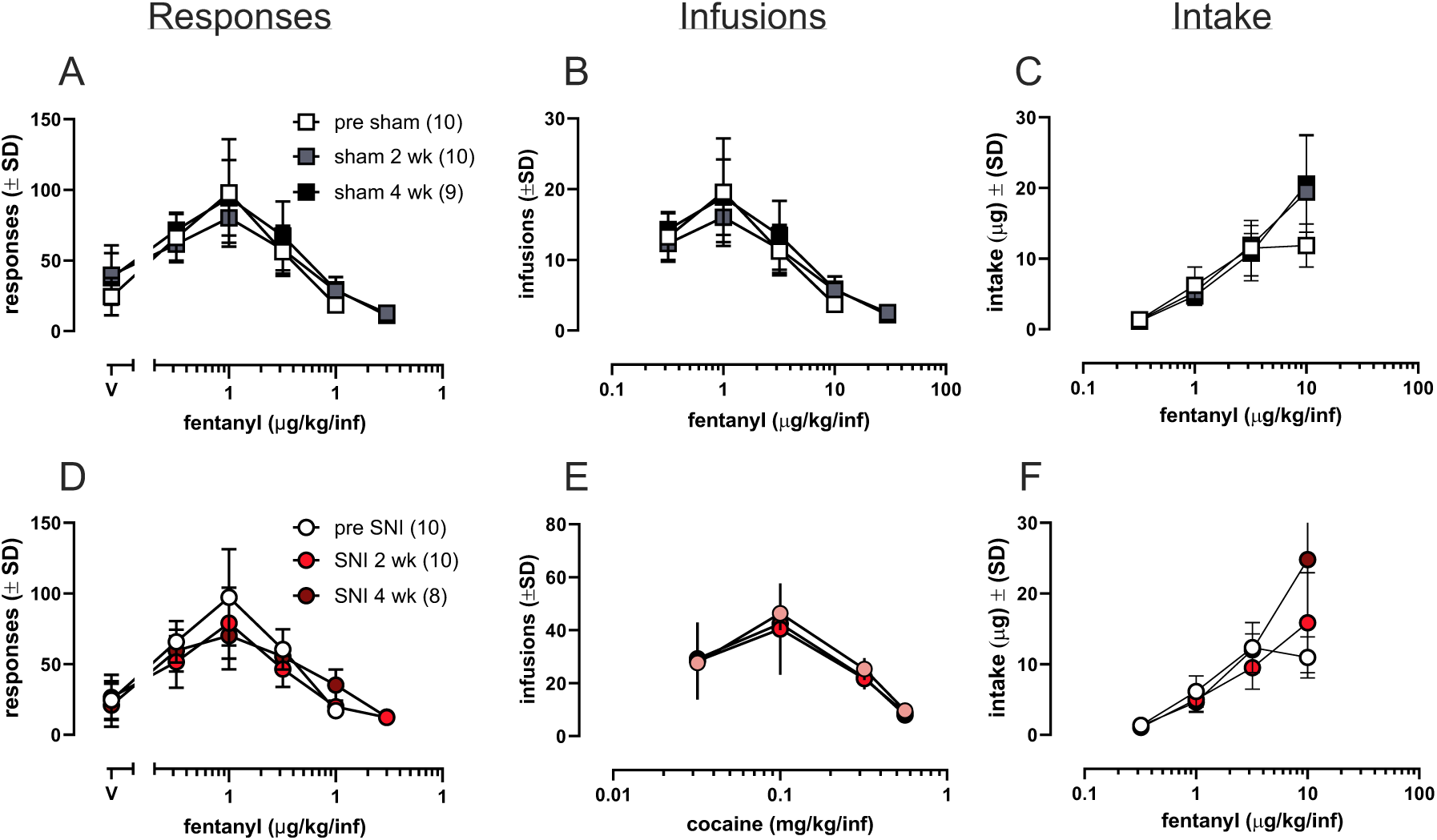
Self-administration of fentanyl prior to and after sham or SNI surgery Data are plotted as responses for fentanyl (A, D), infusions of fentanyl earned (B, E), and total intake (μg) of fentanyl (C, F). Fentanyl maintained self-administration in all rats under a fixed ratio (FR)5 schedule of reinforcement prior to surgery (pre-surgical dose effect curves are the same as plotted in Figure 1). There was no significant difference between fentanyl-maintained responding 2- or 4-weeks following sham (A-C) or SNI (D-F) surgery. After surgery, both sham and SNI groups had increased intake of 10 *u*g/kg fentanyl (C, F). Data are plotted as the mean ± SD. Each data point represents 8-10 rats and include both males and females.

Figure Fig. 2B and E show the number of fentanyl infusions earned for each dose of fentanyl 2- or 4-weeks following surgery. Prior to and after SNI or sham surgery, the number of fentanyl infusions earned per component was dose-dependent, reflected by a main effect of dose [doses included: 0.32-10 ug/kg, F(1.66, 21.62)=68.23, p<0.001, η^2^_p_ =0.84]. There was no difference in fentanyl self-administration between sham and SNI animals suggesting that SNI-induced chronic neuropathic pain did not alter fentanyl self-administration, reflected by a no main effect of surgical status [F(1, 13)=0.41, p=0.46, η^2^_p_ =0.04] (Fig. 2C, F).

There was a significant interaction between surgical status and time, suggesting there were some alterations in number of infusions earned over time in SNI and sham groups [F(2, 26)=4.12, p=0.03, η^2^_p_ =0.24]. This likely reflects small differences between groups over time such as in the SNI group, infusions of 1 *u*g/kg dropped slightly at both post-surgical timepoints as compared with pre-surgery, while in the sham animals, infusions of this dose drop at 2-weeks and returned to pre-surgical levels at 4-weeks. However, there was no three-way interaction between time, dose, and surgical status, suggesting that there were minimal changes in specific doses by surgical status over time [F(6, 78)=1.87, p=0.10, η^2^_p_ =0.13].

Figure 2C and F show the total fentanyl intake for each dose of fentanyl 2- or 4-weeks following surgery. Fentanyl intake was dose dependent, reflected by a main effect of dose [F(4, 52)=189.02, p<0.001, η^2^_p_ =0.94]. While there was no main effect of time [F(1.24, 12.36)=2.85, p=0.11, η^2^_p_ =0.22], there was a significant interaction between time and dose, suggesting intake at specific doses changed over time [F(6, 78)=12.46, p<0.001, η^2^_p_ =0.49]; for example, the intake of 10 *u*g/kg/inf fentanyl increased in the SNI group from the pre-surgical and 2-week timepoints to the 4-week timepoint. Lastly, there was no main effect of surgical status, indicating that fentanyl intake was not significantly different between sham and SNI groups [F(1, 13)=0.27, p=0.61, η^2^_p_ =0.02]. Further, there was no interaction between surgical status and time [F(1.3, 16.89)=0.77, p=0.43, η^2^_p_ =0.06], suggesting that intake of fentanyl was not changed over time by surgical status. All other interaction terms failed to reach significance (p’s>0.12).

### Figure 3. Cocaine self-administration prior to surgery and on post-operative days 4 and 9

Figure 3A and D show the cocaine-maintained responding for each dose cocaine on post-operative days 4 and 9. Cocaine maintained dose-dependent responding, supported by a main effect of dose [F(3, 18)=41.56, p<0.001, η^2^_p_ =0.88]. Though there appeared to be a decrease in responding in both sham (A) and SNI (D) groups on day 4, this was not a significant from pre-surgical levels, supported by no main effect of time [F(1.53, 9.16)=0.19, p=0.78, η^2^_p_ =0.03] as well as no interaction between time and dose [F(6, 36)=0.76, p=0.61, η^2^_p_ =0.11]. Further, there was not no main effect of surgical status [F(1, 6)=0.007, p=0.94, η^2^_p_ =0.001] and no interaction between time and surgical status [F(1.53, 9.16)=0.54, p=0.55, η^2^_p_ =0.08]. Collectively these results demonstrated no significant change in cocaine-maintained responding over time or between SNI and sham groups.

**Figure 3.**
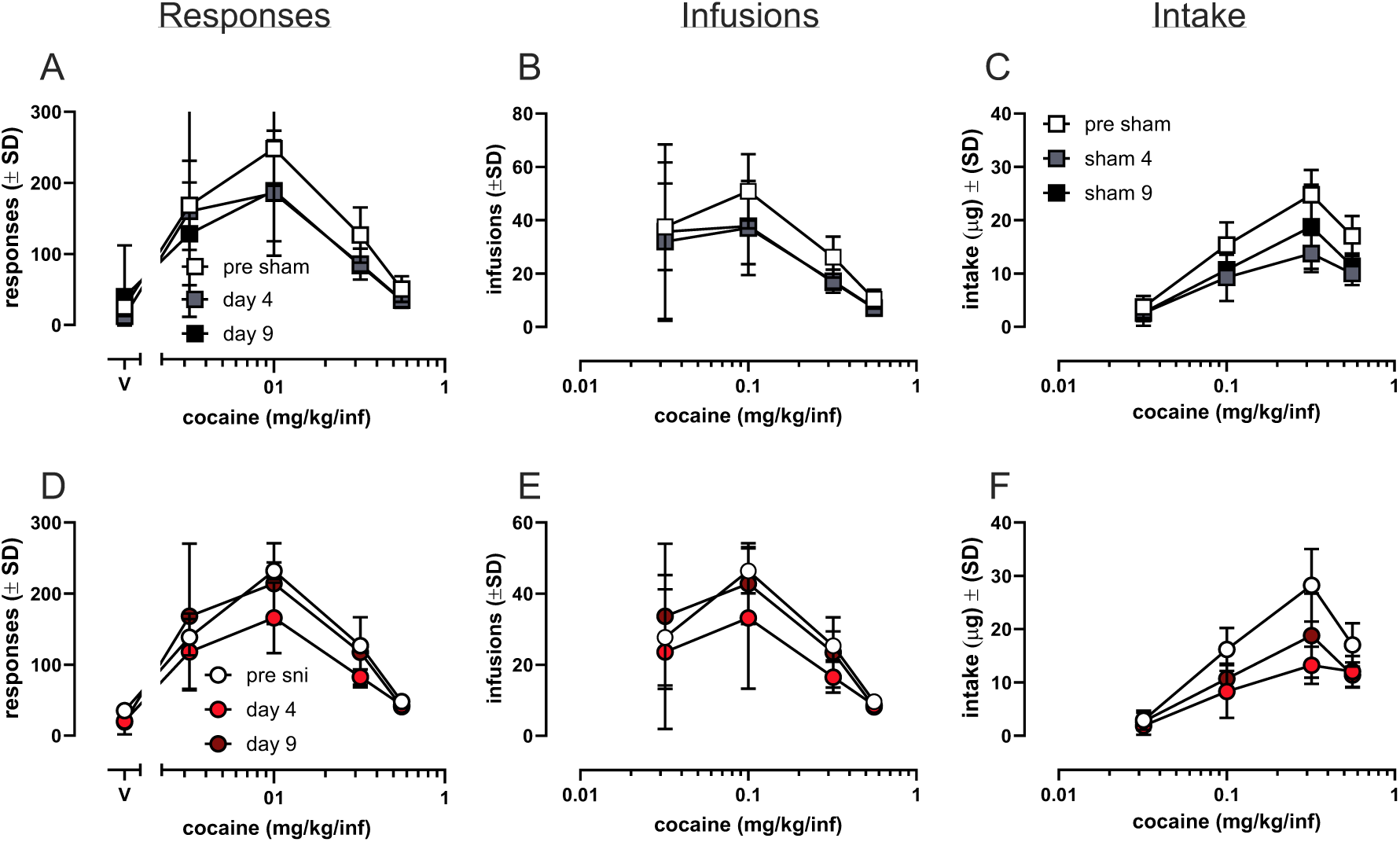
Self-administration of cocaine prior to surgery and on post-operative days 4, 9 Data are plotted as responses for cocaine (A, D), infusions of cocaine earned (B, E), and total intake (μg) of cocaine (C, F). Cocaine maintained self-administration in all rats under a fixed ratio (FR)5 schedule of reinforcement prior to surgery. There was no significant difference in cocaine self-administration on post-operative days 4 and 9 compared to pre-surgical dose effect curves in either sham (A-C) or SNI (D-F) groups. Data are plotted as the mean ± SD. Each data point represents 8-10 rats and include both males and females.

Figure 3B and E show the infusions earned for each dose of fentanyl tested on post-operative days 4 and 9. The number of infusions earned was dose-dependent, supported by a main effect of dose [F(1.65, 9.91)=18.73, p<0.001, η^2^_p_ =0.76]. While there was no main effect of time [F(2, 12)=0.13, p=0.88, η^2^_p_ =0.02], there was a significant interaction between time and dose [F(6, 36)=2.69, p=0.03, η^2^_p_ =0.31], suggesting that over time, there was a dose-dependent change in number of cocaine infusions. This likely reflects the decrease in infusions earned on post-operative day 4 in both sham (B) and SNI (E) groups.

While number of cocaine infusions earned was subject to dose-dependent changes over time, these changes were not explained by surgical status. This was supported by no main effect of time [F(2, 12)=0.47, p=0.64, η^2^_p_ =0.07] and no interaction between time and surgical status [F(2, 12)=0.47, p=0.64, η^2^_p_ =0.07]. Collectively, these results suggest that surgical status did not explain differences in cocaine infusions over time.

Figure 3C and F show cocaine intake for each dose of cocaine on post-operative days 4 and 9. Cocaine intake was dose-dependent, reflected by a main effect of dose [F(1.78, 10.71)=120.08, p<0.001, η^2^_p_ =0.95]. Cocaine intake was subject to changes over time, supported by a main effect of time [F(1.43, 8.60)=7.51, p=0.02, η^2^_p_ =0.56], and these changes were most apparent at larger doses of cocaine (0.1-0.56 mg/kg/inf). This was supported by a significant interaction between time and dose [F(6, 36)=4.58, p=0.002, η^2^_p_ =0.43].

While cocaine intake was subject to changes over time, these changes were not explained by surgical status [no main effect, surgical status, F(1, 6)=0.13, p=0.73, η^2^_p_ =0.02]. this was further supported by no interaction between time and surgical status [F(1.43, 8.60)=0.38, p=0.63, η^2^_p_ =0.06]. Collectively, these results suggest that cocaine intake was subject to small, significant shifts over time that were not explained by surgical status.

### Figure 4. Cocaine self-administration prior to surgery and over 2- or 4-weeks after surgery

Figure 4A and D show cocaine-maintained responding for each dose of cocaine prior to and over 2- or 4-weeks following surgery. Prior to and after surgery, cocaine maintained responding in a dose-dependent manner, reflected by a main effect of dose [doses included: 0.032-0.56 mg/kg F(2.17, 15.17)=80.95, p<0.001, η^2^_p_ =0.92]. Cocaine-maintained responding was not altered by up to 4 weeks of SNI or sham states, reflected by no main effect of time [F(2, 14)=0.01, p=0.99, η^2^_p_ =0.001]. There was also no difference in cocaine-maintained responding between SNI and sham groups, reflected by no main effect of surgical status [F(1, 7)=0.79, p=0.41, η^2^_p_ =0.10]. Further, there was no interaction between time and dose for this dataset [F(8, 56)=0.86, p=0.55, η^2^_p_ =0.11], and there was no interaction between time, dose, and surgical status [F(8, 56)=0.21, p=0.99, η^2^_p_ =0.03]. All other interaction terms failed to reach significance as well (p’s>0.38).

**Figure 4.**
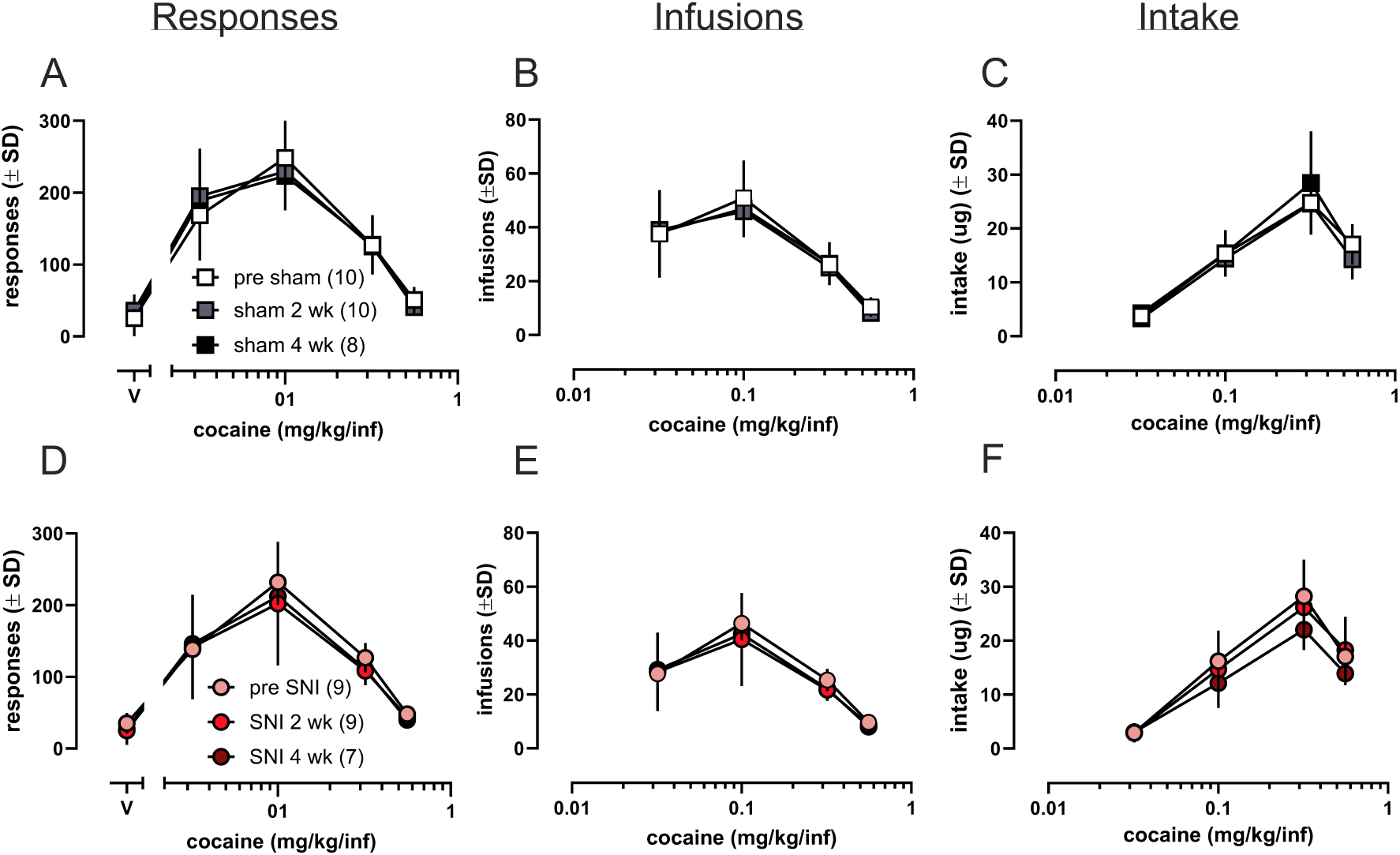
Figure 3-5. Self-administration of cocaine prior to and after surgery Data are plotted as responses for cocaine (A, D), infusions of cocaine earned (B, E), and total intake (μg) of cocaine (C, F). Cocaine maintained self-administration in all rats under a fixed ratio (FR)5 schedule of reinforcement prior to surgery. There was no significant difference in cocaine self-administration 2- or 4-weeks following either sham (A-C) or SNI surgery (D-F). Data are plotted as the mean ± SD. Each data point represents 8-10 rats and include both males and females.

Figure 3-4 B and E show the number of cocaine infusions earned in each component of the multidose procedure 2- or 4-weeks following surgery. The number of cocaine infusions was dose dependent, reflected by a main effect of dose [F(4, 28)=80.95, p<0.001, η^2^_p_ =0.92]. There was no main effect of surgical status suggesting that there was no difference in number of infusions self-administered between sham and SNI groups [no main effect, surgical status: F(1,7)=0.79, p=0.41, η^2^_p_ =0.10]. Further, there was no main effect of time, suggesting cocaine infusions were unaltered over the course of 4 weeks of SNI or sham states [F(2, 14)=0.01, p=0.99, η^2^_p_ =0.001]. There also was not an interaction between time and dose [F(8, 56)=0.86, p=0.55, η^2^_p_=0.11] or time, dose, and surgical status [F(8, 56)=0.21, p=0.99, η^2^_p_ =0.03]. All other interaction terms failed to reach significance (p’s>0.39).

Figure 4C and F show the total cocaine intake in each component of the multidose procedure 2- or 4-weeks following surgery. Prior to surgery, cocaine intake increased as dose increased, through 0.32 mg/kg infusion, while intake decreased at the 0.56 mg/kg/inf dose, reflected by a significant main effect of dose [F(1, 9)=4, 36)=237.19, p<0.001, η^2^_p_ =0.96]. There was no difference in cocaine intake between SNI and sham groups, reflected by no main effect of surgical status [F(1, 9)=0.63, p=0.63, η^2^_p_ =0.03]. Cocaine intake was not altered by up to four weeks of SNI or sham states, reflected by no main effect of time [F(2, 18)=0.042, p=0.96, η^2^_p_ =0.005]. Further, there was no interaction between dose and time [F(8, 72)=0.25, p=0.98, η^2^_p_ =0.03] or dose, time, and surgical status [F(8, 72)=1.61, p=0.14, η^2^_p_ =0.15]. All other interaction terms failed to reach significance (p’s>0.14).

### Figure 5. Antihyperalgesic- and antinociceptive-like effects of intravenous fentanyl or cocaine in the Randall Selitto Assay

The data shown in Figure 5A and B demonstrate that i.v. fentanyl produced dose-dependent antinociceptive-like and antihyperalgesic-like effects on days 4 and 9 following surgery. We evaluated 0.32-32 μg/kg fentanyl as these are the doses available following surgery. SNI surgery induced a hyperalgesic-like state, as demonstrated by lowered baseline paw withdrawal thresholds, reflected by a main effect of surgical status [F(1, 20)=2.97, p<0.001, η^2^_p_ =0.65]. There was a significant main effect of dose such that fentanyl produced dose dependent increases in paw withdrawal thresholds [F(2.9, 58.07)=495.69, p<0.001, η^2^_p_ =0.96]. Additionally, there was a dose by surgical status interaction [F(2.90, 58.07)=19.76, p<0.001, η^2^_p_ =0.50]. This reflects lower baseline paw withdrawal thresholds in the SNI group with no fentanyl on board and low dose fentanyl (0.32 and 1 *u*g/kg). A one-way ANOVA split by surgical status and Tukey’s post hoc analyses was used to determine which doses of fentanyl significantly increased paw withdrawal thresholds compared to vehicle in both sham and SNI groups. There was a main effect of dose in both sham [F(5, 66)=116.73, p<0.001, η^2^_p_ =0.90] and SNI groups [F(5, 66)=212.84, p<0.001, η^2^_p_ =0.94]. In sham rats, 3.2-32 *u*g/kg fentanyl significantly increased paw withdrawal thresholds as compared with vehicle (all p<0.001). In SNI rats, 1 (p=0.003), 3.2 (p<0.001), 10 (p<0.001), and 32 (p<0.001) *u*g/kg fentanyl significantly increased paw withdrawal thresholds as compared with vehicle.

**Figure 5.**
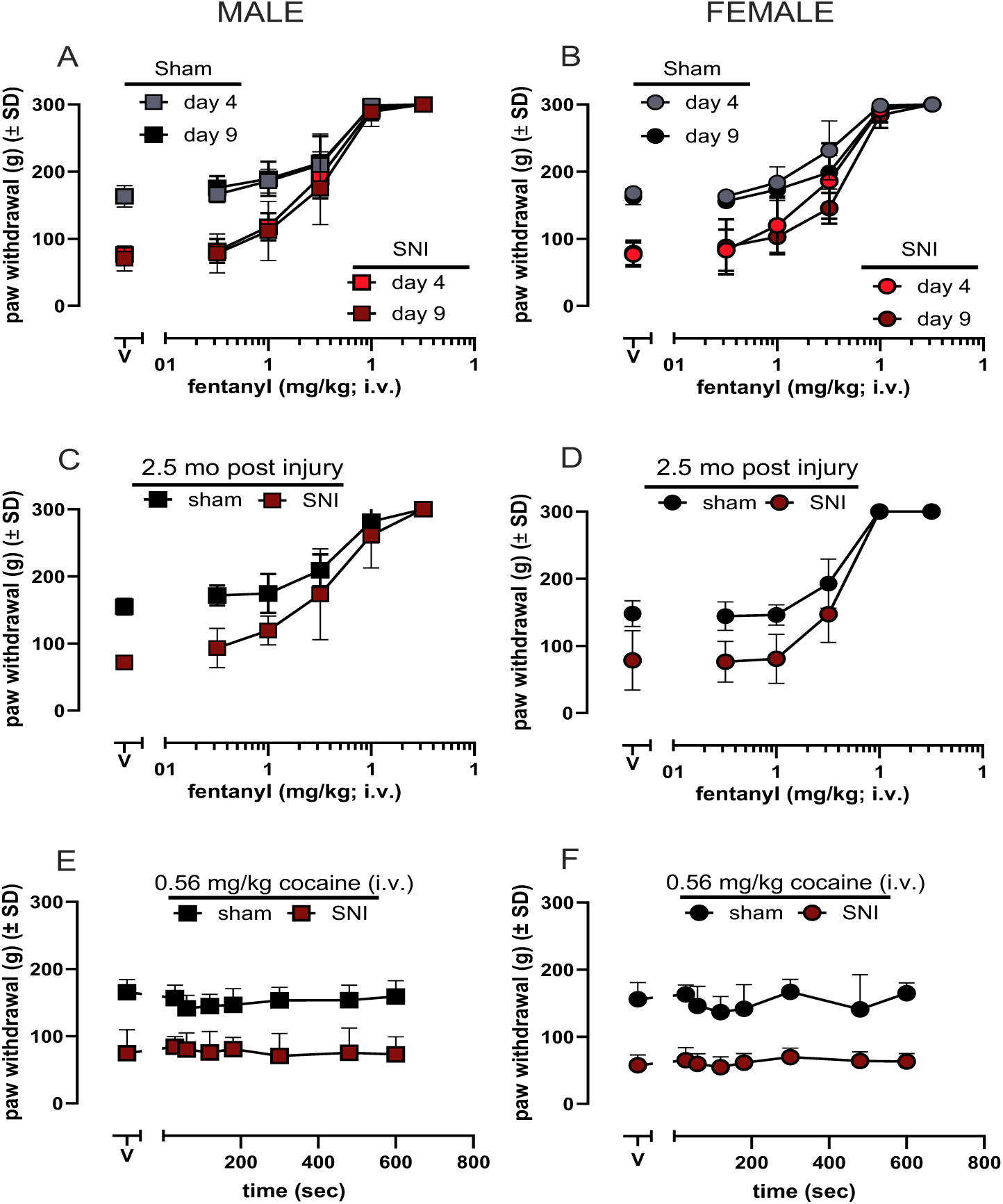
Effect of intravenous fentanyl or cocaine on paw withdrawal thresholds Effects of intravenous fentanyl (0-32 μg/kg) or cocaine (0.56 mg/kg) on paw withdrawal thresholds in male and female rats. Fentanyl induced antinociceptive- and antihyperalgesic-like effects were evaluated on post-operative days 4 and 9 (A, B) as well as 2.5 months following surgery (C, D). Cocaine-induced alterations in paw withdrawal thresholds were evaluated once, as a time course (E, F). Data are plotted as the mean ± SD paw withdrawal threshold. Each datapoint represents the average of 6 rats.

Fentanyl was approximately equally effective on day 4 and 9 following surgery, as reflected by a non-significant main effect of time [F(1, 20)=1.19, p=0.29, η^2^_p_ =0.06] as well as a no interaction between time and dose [F(5, 100)=1.63, p=0.16, η^2^_p_ =0.08]. There was no sex difference in fentanyl-induced antinociceptive- or antihyperalgesic-like effects [F(1, 20)=0.08, p=0.78, η^2^_p_ =0.004]. All other interaction terms failed to reach significance (p’s<0.1).

As shown in Figure 5C and D, i.v. fentanyl produced dose dependent antinociceptive- and antihyperalgesic-like effects 2.5 months after injury. SNI surgery induced a hyperalgesic-like state as demonstrated by lowered baseline paw withdrawal thresholds as compared with sham, reflected by a main effect of surgical status [F(1, 20)=51.62, p<0.001, η^2^_p_ =0.72]. There was a significant main effect of dose such that fentanyl produced dose dependent increases in paw withdrawal thresholds [F(2.88, 57.63)=278.03, p<0.001, η^2^_p_ =0.93]. A one-way ANOVA split by surgical status and Tukey’s post hoc analyses was used to determine which doses of fentanyl significantly increased paw withdrawal thresholds compared to vehicle in both sham and SNI groups. There was a significant main effect of dose in sham groups [F(5, 66)=118.67, p<0.001, η^2^_p_ =0.9] and SNI groups [F(5, 66)=105.52, p<0.001, η^2^_p_ =0.89]. 3.2-32 *u*g/kg fentanyl were significantly more effective than vehicle in sham (all p<0.001) and SNI groups (all p<0.001).

There was no main effect of sex [main effect: sex, [F(1, 20)=1.64, p=0.22, η^2^_p_ =0.08]. Though, there was an interaction between dose and sex, suggesting differences in the fentanyl dose effect curves by sex [F(2.88, 57.63)=4.88, p=0.005, η^2^_p_ =0.20]. This likely represents the slightly increased effectiveness of 1 and 3.2 *u*g/kg in males (Fig. 5C). All other interactions failed to reach significance, including the three-way interaction term (p’s>0.86).

We tested 0.56 mg/kg cocaine as it is the highest dose available in self-administration assays, and 0.56 mg/kg was tested as a time course. As shown in Figure 5E and F, i.v. cocaine (0.56 mg/kg) failed to attenuate SNI-induced mechanical hypersensitivity in either sex. 0.56 mg/kg cocaine (i.v.) failed to alter paw withdrawal over a 10 min period in males (Fig. 5E) and females (Fig. 5F), reflected by a non-significant main effect of time [F(4.12, 82.33)=2.04, p=0.10, η^2^_p_ =0.09]. There was significant difference between sham and SNI groups such that, at all points tested, SNI groups had lower paw withdrawal thresholds than sham groups [main effect: surgical status, F(1, 20)=132.82, p<0.001, η^2^_p_ =0.87]. Lastly, there was no main effect of sex [main effect: sex, F(1, 20)=1.16, p=0.29, η^2^_p_ =0.06], indicating a lack of sex differences. All interaction terms failed to reach significance (p’s>0.17)

## Discussion

The goal of the present study was to evaluate if SNI-induced hypersensitivity altered ongoing self-administration of fentanyl or cocaine. Therefore, animals were first trained to self-administer fentanyl *or* cocaine, then underwent SNI or sham surgery, were allowed 72 hours of recovery before resuming self-administration on post-operative day 4, and post-operative evaluation of fentanyl-maintained behavior continued for 4 weeks. At all timepoints, the dose effect curves were examined across three variables (i.e., responding, infusions, intake). Prior to surgery, fentanyl-maintained responding and intake dose-dependently increased (Fig. 1). Compared with fentanyl dose effect curves established prior to surgery, fentanyl-maintained behavior was significantly decreased on post-operative day 4 in both sham and SNI groups (Fig. 1A, D). However, this decrease in fentanyl self-administration did not persist over the remaining duration of post-operative evaluation. By postoperative day 9, fentanyl self-administration behaviors returned to approximately pre-surgical levels in both sham and SNI groups (Fig. 1). Decreases in fentanyl self-administration were observed in both sham and SNI groups, suggesting SNI-induced hypersensitivity does *not* explain the observed decreases in fentanyl self-administration. While sham surgery does not include damage to the sciatic nerve, a large muscle incision is made. Although the sham groups received carprofen to manage post-operative pain and/or discomfort, post-operative pain or surgery itself and exposure to anesthesia may contribute to the decrease in fentanyl self-administration immediately after surgery. Decreased self-administration in sham groups has been observed before in other studies in mice (Wade et al., 2003).

To evaluate if fentanyl effectively alleviates SNI-induced hypersensitivity to mechanical stimulation on postoperative day 4, a separate group of rats was used to evaluate the effects of intravenous fentanyl on paw withdrawal thresholds. Paw withdrawal thresholds following vehicle administration were similar to thresholds measured prior to surgery. Together, these data suggest that the sham groups were not hypersensitive to mechanical stimulation 4- or 9-days post-surgery. However, it is somewhat challenging to completely dismiss post-operative pain as an explanation for the initial decrease in fentanyl maintained-responding as there is a large difference in the environment between Randall Selitto assays (i.e., animals suspended from hammock-like restraint with no weight on hind paws) and the self-administration chambers (i.e., animals support full body weight and move on grid floors).

Fentanyl self-administration behaviors were examined over a total of 4 weeks following surgery in order to determine if chronic pain states alter the ongoing self-administration of MOR agonists over an extended period of time. The initial decrease in fentanyl self-administration was not persistent, such that, at 2- or 4-weeks following surgery, there were no robust shifts in the fentanyl dose effect curves. Nevertheless, there was evidence of increased intake of specifically 10 ug/kg fentanyl in both SNI and sham groups (Fig. 3). This finding is consistent with decreased sensitivity, or tolerance development, to MOR agonist-induced effects following repeated self-administration (Dao et al., 2021; Malone et al., 2021).

It is also possible that tolerance developed to the rate suppressant effects of fentanyl (rather than the reinforcing effects), such that animals experience less behavioral disruption or sedation at larger doses over time and, therefore, intake of large fentanyl doses could increase. Repeated administration of MOR agonists has been previously demonstrated to produce rightward shifts in the opioid analgesic-induced locomotor suppressing effects (Timar, Gyarmati, & Furst, 2005). Overall, in the present study, while there were changes in fentanyl intake over time, SNI-induced hypersensitivity failed to directly alter the ongoing self-administration of fentanyl. This finding is consistent with several previous studies in which acute or sub-chronic pain states did not alter ongoing opioid self-administration (Barattini et al., 2023; Reiner et al., 2021).

However, several other studies report an increase or decrease in the self-administration of MOR agonists in the presence of a pain state. While there are many methodological differences in the present and previous studies, two factors may contribute to the differential effects of pain on opioid self-administration across studies: (1) when pain states were induced (prior to or after acquisition of self-administration, and (2) the type and duration of pain used. A large number of the previous studies have induced pain states prior to the acquisition of self-administration (Colpaert et al., 2001; Martin et al., 2007; Higginbotham et al., 2022; Hipolitto et al., 2015; Hou et al., 2015; Lyness et al., 1989; Wade et al., 2003; Woller et al., 2014), while relatively few have induced pain states after acquisition of self-administration (Barattini et al., 2023; Neelakantan et al., 2016; Reiner et al., 2021). Generally, the studies that induced pain states after acquisition do not report an effect of pain states on self-administration behaviors (Barattini et al., 2023; Neelakantan et al., 2016; Reiner et al, 2021), which is consistent with the present study. Only one previous study has examined if chronic neuropathic pain alters ongoing self-administration of MOR agonists, and this study demonstrated that in male mice, paclitaxel-induced hypersensitivity prevented the decrease in morphine breakpoint over time, compared to the sham group (Neelakantan et al., 2016). These findings suggest a very small impact of paclitaxel on morphine-induced reinforcing effects. It is also possible that the type of pain utilized (e.g., neuropathic or inflammatory) may differentially alter the reinforcing effects of MOR agonists. There have been fewer studies assessing the impact of chronic neuropathic pain on opioid self-administration than inflammatory or nociceptive pain states, and the results from the inflammatory or nociceptive pain state studies are quite heterogeneous (Decrease-Wade et al., 2002; Lyness et al., 1989; No change-Barattini et al., 2023; Hou et al., 2015; Reiner et al., 2021; Increase-Colpaert et al., 2001; Dib & Duclaux, 1982; Hipolitto et al., 2015).

Overall, the primary goal of the present study was to determine if chronic neuropathic pain or sham states altered the ongoing self-administration of fentanyl, but we also evaluated if pain states alter the reinforcing effects of a non-opioidergic drug of abuse-cocaine, which no previous studies have directly tested. Collectively, the present study demonstrated that SNI-induced hypersensitivity did not alter the reinforcing effects of fentanyl or cocaine. Though, we observed changes in fentanyl induced reinforcing effects over time, such that animals in both sham and SNI groups responded more for 10 *u*g/kg/inf fentanyl, SNI-induced hypersensitivity did not explain the increased intake of 10 *u*g/kg/inf fentanyl.

The failure of SNI-induced hypersensitivity to alter reinforcing effects or antinociceptive- and antihyperalgesic-like effects of fentanyl could be due to use of a high efficacy MOR agonist in these experiments. Previous studies have reported that pain states generally reduce MOR expression and activity in reward circuitry (Back et al., 2006; Campos-Jurado et al, 2019; Dong et al., 2019; Hou et al., 2017; Ji et al., 1995; Kaneuchi et al., 2019; Ozaki et al., 2002; Pol et al., 2006; Porecca et al., 1998; Thompson et al., 2018; Yamamoto et al., 2008; Zhang et al., 1998); however, it is possible these changes in receptor expression or activity may not be sufficient to alter behavioral effects of a high efficacy agonist. Future studies should examine the reinforcing effects of a lower efficacy MOR agonist such as nalbuphine or buprenorphine and the extent to which chronic pain alters learning.

## Author Contributions

Participated in research design: Burgess, Traynor, Jutkiewicz

Conducted experiments: Burgess

Performed data analysis: Burgess, Jutkiewicz

Wrote or contributed to the writing of the manuscript: Burgess, Traynor, Jutkiewicz

## Footnotes

This work was supported by the Dr. Ben & Diana Lucchesi Graduate Education Fellowship (GEB); and the National Institutes of Health National Institute on Drug Abuse [Grants T32 DA007281 (GEB), UG3 DA056884 (JRT)].

## Bibliography

1. Hser, Y. I., Mooney, L. J., Saxon, A. J., Miotto, K., Bell, D. S., & Huang, D. (2017). Chronic pain among patients with opioid use disorder: results from electronic health records data. Journal of substance abuse treatment, 77, 26–30.

2. Latif, Z. E. H., Skjærvø, I., Solli, K. K., & Tanum, L. (2021). Chronic pain among patients with an opioid use disorder. The American journal on addictions, 30(4), 366–375.

3. Hoffman, E. M., Watson, J. C., St Sauver, J., Staff, N. P., & Klein, C. J. (2017). Association of long-term opioid therapy with functional status, adverse outcomes, and mortality among patients with polyneuropathy. JAMA neurology, 74(7), 773–779.

4. Baumann, L., Bello, C., Georg, F. M., Urman, R. D., Luedi, M. M., & Andereggen, L. (2023). Acute Pain and Development of Opioid Use Disorder: Patient Risk Factors. Current pain and headache reports, 27(9), 437–444.

5. Brummett, C. M., Waljee, J. F., Goesling, J., Moser, S., Lin, P., Englesbe, M. J., Bohner, A. S. B., Kheterpal, S., & Nallamothu, B. K. (2017). New persistent opioid use after minor and major surgical procedures in US adults. JAMA surgery, 152(6), e170504–e170504.

6. Katz, C., El-Gabalawy, R., Keyes, K. M., Martins, S. S., & Sareen, J. (2013). Risk factors for incident nonmedical prescription opioid use and abuse and dependence: results from a longitudinal nationally representative sample. Drug and alcohol dependence, 132(1-2), 107–113.

7. Cahill, C. M., Xue, L., Grenier, P., Magnussen, C., Lecour, S., & Olmstead, M. C. (2013). Changes in morphine reward in a model of neuropathic pain. Behavioural pharmacology, 24(3), 207–213.

8. Lim, G., Kim, H., McCabe, M. F., Chou, C. W., Wang, S., Chen, L. L., John, J.A., Marota, A. B., Breiter, H. C., & Mao, J. (2014). A leptin-mediated central mechanism in analgesia-enhanced opioid reward in rats. Journal of Neuroscience, 34(29), 9779–9788.

9. Navratilova, E., Nation, K., Remeniuk, B., Neugebauer, V., Bannister, K., Dickenson, A. H., & Porreca, F. (2020). Selective modulation of tonic aversive qualities of neuropathic pain by morphine in the central nucleus of the amygdala requires endogenous opioid signaling in the anterior cingulate cortex. Pain, 161(3), 609.

10. Zhang, Z., Tao, W., Hou, Y. Y., Wang, W., Lu, Y. G., & Pan, Z. Z. (2014). Persistent pain facilitates response to morphine reward by downregulation of central amygdala GABAergic function. Neuropsychopharmacology, 39(9), 2263–2271.

11. Colpaert, F. C., Tarayre, J. P., Alliaga, M., Slot, L. B., Attal, N., & Koek, W. (2001). Opiate self-administration as a measure of chronic nociceptive pain in arthritic rats. Pain, 91(1-2), 33–45.

12. Higginbotham, J. A., Abt, J. G., Tiech, R. H., & Morón, J. A. (2022). Time-dependent enhancement in ventral tegmental area dopamine neuron activity drives pain-facilitated fentanyl intake in males. bioRxiv, 2022–08.

13. Hipólito, L., Wilson-Poe, A., Campos-Jurado, Y., Zhong, E., Gonzalez-Romero, J., Virag, L., Whittington, R., Comer, S.D., Carlton, S. M., Walker, B. M., Bruchas, M. R., & Morón, J. A. (2015). Inflammatory pain promotes increased opioid self-administration: role of dysregulated ventral tegmental area μ opioid receptors. Journal of Neuroscience, 35(35), 12217–12231.

14. Ozaki, S., Narita, M., Narita, M., Iino, M., Sugita, J., Matsumura, Y., & Suzuki, T. (2002). Suppression of the morphine-induced rewarding effect in the rat with neuropathic pain: implication of the reduction in µ-opioid receptor functions in the ventral tegmental area. Journal of neurochemistry, 82(5), 1192–1198.

15. Suzuki, T., Kishimoto, Y., & Misawa, M. (1996). Formalin-and carrageenan-induced inflammation attenuates place preferences produced by morphine, methamphetamine and cocaine. Life sciences, 59(19), 1667–1674.

16. Nazarian, A., Negus, S. S., & Martin, T. J. (2021). Factors mediating pain-related risk for opioid use disorder. Neuropharmacology, 186, 108476.

17. Narita, M., Iino, M., Sugita, J., Matsumura, Y., & Suzuki, T. (2002). Suppression of the morphine-induced rewarding effect in the rat with neuropathic pain: implication of the reduction in µ-opioid receptor functions in the ventral tegmental area. Journal of neurochemistry, 82(5), 1192–1198.

18. Wade, C. L., Krumenacher, P., Kitto, K. F., Peterson, C. D., Wilcox, G. L., & Fairbanks, C. A. (2013). Effect of chronic pain on fentanyl self-administration in mice. PLoS One, 8(11), e79239.

19. Martin, T. J., Kim, S. A., Buechler, N. L., Porreca, F., & Eisenach, J. C. (2007). Opioid self-administration in the nerve-injured rat: relevance of antiallodynic effects to drug consumption and effects of intrathecal analgesics. The Journal of the American Society of Anesthesiologists, 106(2), 312–322

20. Barattini, A. E., Montanari, C., Edwards, K. N., Edwards, S., Gilpin, N. W., & Pahng, A. R. (2023). Chronic inflammatory pain promotes place preference for fentanyl in male rats but does not change fentanyl self-administration in male and female rats. Neuropharmacology, 231, 109512.

21. Hou, Y. Y., Cai, Y. Q., & Pan, Z. Z. (2015). Persistent pain maintains morphine-seeking behavior after morphine withdrawal through reduced MeCP2 repression of GluA1 in rat central amygdala. Journal of Neuroscience, 35(8), 3689–3700.

22. Reiner, D. J., Townsend, E. A., Orihuel, J., Applebey, S. V., Claypool, S. M., Banks, M. L., Shaham, Y., & Negus, S. S. (2021). Lack of effect of different pain-related manipulations on opioid self-administration, reinstatement of opioid seeking, and opioid choice in rats. Psychopharmacology, 238, 1885–1897.

23. Shippenberg, T. S., Emmett-Oglesby, M. W., Ayesta, F. J., & Herz, A. (1988). Tolerance and selective cross-tolerance to the motivational effects of opioids. Psychopharmacology, 96, 110–115.

24. Neelakantan, H., Ward, S. J., & Walker, E. A. (2016). Effects of paclitaxel on mechanical sensitivity and morphine reward in male and female C57Bl6 mice. Experimental and clinical psychopharmacology, 24(6), 485.

25. Decosterd, I., & Woolf, C. J. (2000). Spared nerve injury: an animal model of persistent peripheral neuropathic pain. Pain, 87(2), 149–158.

26. Dao, A. N., Beacher, N. J., Mayr, V., Montemarano, A., Hammer, S., & West, M. O. (2021). Chronic fentanyl self-administration generates a shift toward negative affect in rats during drug use. Brain Sciences, 11(8), 1064.

27. Malone, S. G., Keller, P. S., Hammerslag, L. R., & Bardo, M. T. (2021). Escalation and reinstatement of fentanyl self-administration in male and female rats. Psychopharmacology, 238(8), 2261–2273.

28. Picetti, R., Caccavo, J. A., Ho, A., & Kreek, M. J. (2012). Dose escalation and dose preference in extended-access heroin self-administration in Lewis and Fischer rats. Psychopharmacology, 220, 163–172.

29. Timár, J., Gyarmati, Z., & Fürst, Z. (2005). The development of tolerance to locomotor effects of morphine and the effect of various opioid receptor antagonists in rats chronically treated with morphine. Brain research bulletin, 64(5), 417–424.

30. Lyness, W. H., Smith, F. L., Heavner, J. E., Iacono, C. U., & Garvin, R. D. (1989). Morphine self-administration in the rat during adjuvant-induced arthritis. Life sciences, 45(23), 2217–2224.

31. Woller, S. A., Malik, J. S., Aceves, M., & Hook, M. A. (2014). Morphine self-administration following spinal cord injury. Journal of Neurotrauma, 31(18), 1570–1583.

32. Back, S. K., Lee, J., Hong, S. K., & Na, H. S. (2006). Loss of spinal μ-opioid receptor is associated with mechanical allodynia in a rat model of peripheral neuropathy. Pain, 123(1-2), 117–126.

33. Campos-Jurado, Y., Igual-López, M., Padilla, F., Zornoza, T., Granero, L., Polache, A., Agustín-Pavón, C. & Hipólito, L. (2019). Activation of MORs in the VTA induces changes on cFos expression in different projecting regions: Effect of inflammatory pain. Neurochemistry International, 131, 104521.

34. Dong, J., Zuo, Z., Yan, W., Liu, W., Zheng, Q., & Liu, X. (2019). Berberine ameliorates diabetic neuropathic pain in a rat model: involvement of oxidative stress, inflammation, and μ-opioid receptors. Naunyn-Schmiedeberg’s Archives of Pharmacology, 392, 1141–1149.

35. Hou, X., Weng, Y., Ouyang, B., Ding, Z., Song, Z., Zou, W., … & Guo, Q. (2017). HDAC inhibitor TSA ameliorates mechanical hypersensitivity and potentiates analgesic effect of morphine in a rat model of bone cancer pain by restoring μ-opioid receptor in spinal cord. Brain Research, 1669, 97–105.

36. Ji, R. R., Zhang, Q., Law, P. Y., Low, H. H., Elde, R., & Hokfelt, T. (1995). Expression of mu-, delta-, and kappa-opioid receptor-like immunoreactivities in rat dorsal root ganglia after carrageenan-induced inflammation. Journal of Neuroscience, 15(12), 8156–8166.

37. Kaneuchi, Y., Sekiguchi, M., Kameda, T., Kobayashi, Y., & Konno, S. I. (2019). Temporal and spatial changes of μ-opioid receptors in the brain, spinal cord and dorsal root ganglion in a rat lumbar disc herniation model. Spine, 44(2), 85–95.

38. Ozaki, S., Narita, M., Narita, M., Iino, M., Sugita, J., Matsumura, Y., & Suzuki, T. (2002). Suppression of the morphine-induced rewarding effect in the rat with neuropathic pain: implication of the reduction in µ-opioid receptor functions in the ventral tegmental area. Journal of neurochemistry, 82(5), 1192–1198.

39. Pol, O., Murtra, P., Caracuel, L., Valverde, O., Puig, M. M., & Maldonado, R. (2006). Expression of opioid receptors and c-fos in CB1 knockout mice exposed to neuropathic pain. Neuropharmacology, 50(1), 123–132.

40. Porreca, F., Tang, Q., Bian, D., Riedl, M., Elde, R., & Lai, J. (1998). Spinal opioid mu receptor expression in lumbar spinal cord of rats following nerve injury. Brain research, 795(1-2), 197–203.

41. Thompson, S. J., Pitcher, M. H., Stone, L. S., Tarum, F., Niu, G., Chen, X., Kiesewetter, D.O., Schweinhardt, P., & Bushnell, M. C. (2018). Chronic neuropathic pain reduces opioid receptor availability with associated anhedonia in rat. Pain, 159(9), 1856.

42. Yamamoto, J., Kawamata, T., Niiyama, Y., Omote, K., & Namiki, A. (2008). Down-regulation of mu opioid receptor expression within distinct subpopulations of dorsal root ganglion neurons in a murine model of bone cancer pain. Neuroscience, 151(3), 843–853.

43. Zhang, Q., Schäfer, M., Elde, R., & Stein, C. (1998). Effects of neurotoxins and hindpaw inflammation on opioid receptor immunoreactivities in dorsal root ganglia. Neuroscience, 85(1), 281–291.

